# Neural Evidence of Early Sensitivity to Text in Pre-reading Toddlers

**DOI:** 10.64898/2026.02.24.707347

**Authors:** Nadeen Kherbawy, Chritine E Potter, Sagi Jaffe-Dax

## Abstract

Learning to read leads to widespread changes in brain organization, but it is not yet known when text first becomes a privileged stimulus. To test whether neural sensitivity to text can appear prior to reading instruction, 31 monolingual toddlers in Israel (2.1-3.6 years) not yet enrolled in school were presented with displays of real, native text and visually matched non-text symbols. Using functional near-infrared spectroscopy, we found different patterns of activity in response to text vs. non-text across multiple cortical regions. Most notably, text elicited more activity in the ventrolateral prefrontal cortex, a region often associated with language processing. These results suggest that through implicit sensitivity to regularities in their input, toddlers may be able to discover that text is a meaningful stimulus and could begin to develop associations that will allow them to build connections between language and text.

**Research Highlights:**

⍰ Toddlers show different neural responses to real text vs. non-text symbols.
⍰ Unfamiliar symbols evoke a novelty response in multiple cortical regions.
⍰ Text elicits more activity in a left ventrolateral prefrontal cortex, a region associated with processing language.
⍰ Before they know how to read, toddlers may recognize text as a frequent, familiar stimulus that is linked to language.

## Introduction

Reading provides a foundation for academic achievement, and children who read well early tend to succeed in school and beyond (Hernandez, 2011; Rabiner et al., 2016). Neuroimaging studies show that patterns of young children’s brain activation during reading and language processing predict reading proficiency (McNorgan et al., 2011), and gains in reading skill are reflected in increasingly specialized neural responses to text (Cantlon et al., 2011; Dundas et al., 2012; Wang et al., 2024). Pinpointing the neural substrates involved in early reading-related processing is therefore essential for understanding the developmental trajectory of reading and potentially identifying children at risk for reading impairments, enabling earlier and targeted interventions. This study investigated whether the beginnings of neural sensitivity to text can be observed in pre-reading toddlers, thereby providing insight into the emergence of the neural architecture that supports reading.

Reading alters cognition and the brain, especially for those that learn to read as children (Dehaene et al., 2010; Thiebaut de Schotten et al., 2014). Text becomes privileged, such that proficient readers struggle to suppress the automaticity of reading even when it interferes with task goals (Stroop, 1935). The visual word form area (VWFA), a cortical area within the left fusiform gyrus, has gained attention as a key region linked to reading (Dehaene et al., 2010). Additionally, although text is a visual stimulus, it evokes selective responses in cortical regions that constitute a subset of the language processing network, including the superior temporal cortex (STC), ventrolateral prefrontal cortex (VLPFC), and dorsal premotor cortex (pMD) in the left hemisphere (Dehaene et al., 2010; Fedorenko et al., 2011; see Figure 1). These regions are thought to facilitate associations between visual input and linguistic representations (Cohen et al., 2002). Thus, activity can provide evidence that a viewer not only recognizes text but also has formed links between its visual form and communicative function.

**Figure 1.**
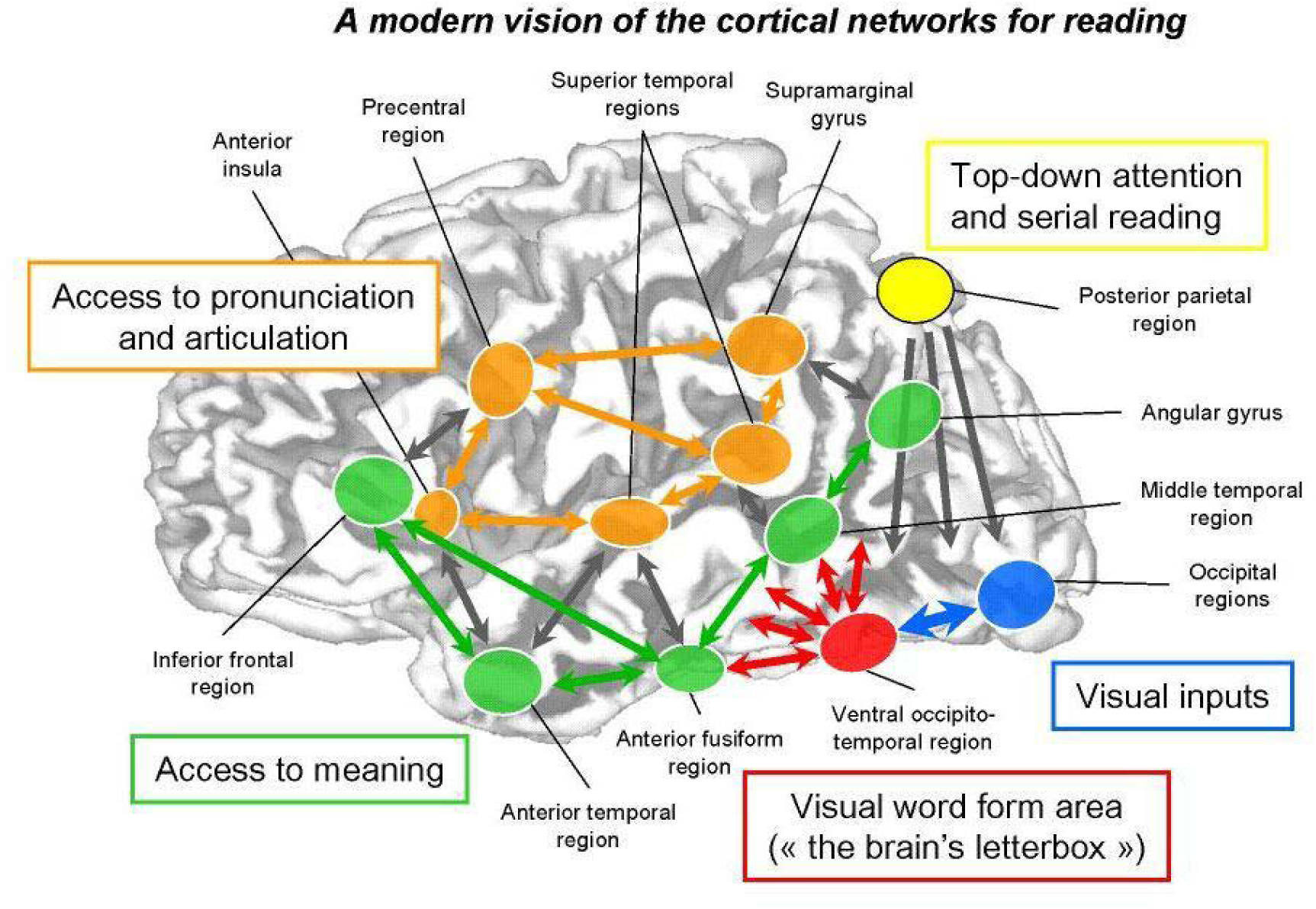
The Reading Network. *Note*. Illustration of the brain regions suggested to be involved in skilled reading and the interactions between them. Reproduced with permission from Dehaene, 2009, p. 4.

Importantly, learning to read is a gradual process. Evidence from four- to five-year-old children learning to read suggests the selectivity of the reading network emerges slowly (Cantlon et al., 2011; Saygin et al., 2016; Schlaggar & McCandliss, 2007). Even as children begin to read more fluently, they exhibit less selective neural responses to text compared to adults (Cantlon et al., 2011; Crollen et al., 2025; Schlaggar & McCandliss, 2007; Zhang et al., 2024). Reading-related activities are associated with neural activity across a wide range of regions for early readers, and as children’s reading skills improve, they show more consistent patterns (Jangraw et al., 2023; Pugh et al., 2013; Smith et al., 2018; Wat et al., 2024; Yamada et al., 2011). Moreover, longitudinal findings indicate that improvements in reading proficiency are accompanied by domain-general changes in brain organization, underscoring the bidirectional relationship between reading and neural development (Centanni et al., 2018; Dundas et al., 2013).

While many studies have documented influences of learning to read on brain organization (Cantlon et al., 2011; Dundas et al., 2013; Wang et al., 2022), no study to date has investigated neural origins of text sensitivity in children prior to formal reading instruction. Behavioral measures of early literacy skills (e.g., familiarity with sounds or letter names) are evident in children as young as two years and predict later reading skills (Kenner et al., 2017; Kirby et al., 2003). In addition, neural activity associated with sound- and language-based activities during the preschool years is correlated with children’s subsequent reading skills (Jasińska et al., 2021). One explanation for the links between pre-reading abilities and eventual reading proficiency is that some children have more experience with literacy activities (Yeo et al., 2014) and therefore more exposure to text and its co-occurrence with dense language input (Montag et al., 2015). Infants’ attention is drawn to frequent stimuli (Amso & Scerif, 2015), and cortical regions develop specialized responses to specific types of input in tandem with increased attention to that type of input (Johnson, 2000). Thus, regular exposure to text could establish the neural foundations for later reading development. We tested the hypothesis that toddlers recognize text as a meaningful stimulus and have begun to form associations between text and spoken language, reflected in different neural responses to text compared to other stimuli. That is, we propose that neural changes supporting reading emerge at early stages of exposure to text.

To test this proposal, we used functional near-infrared spectroscopy (fNIRS), an infant-friendly, non-invasive imaging technique that is less susceptible to movement artifacts while monitoring cortical function by measuring changes in blood oxygenation (Lloyd-Fox et al., 2010). Researchers are beginning to use fNIRS to measure reading-related brain activity in children, and there is promising evidence that it will be useful in revealing developmental changes (see Bode et al., 2026 for review). We measured the cortical activity of two-to three-year-old toddlers while they viewed familiar text or non-text symbols. We focused on the VLPFC, STC, pMD, and parietal regions of the reading network due to their role in facilitating connections between visual input and language in the language network (Fedorenko et al., 2011). We did not include other regions, such as the VWFA or the bilateral ventral occipital regions, because they cannot be reliably measured using fNIRS (Chen et al., 2020) and because our question was whether toddlers had begun to form associations between text and language. We predicted toddlers would show greater activity in response to text, reflecting their prioritization of a familiar stimulus and that activity would be left lateralized, indicating an early link between text and language. Children growing up in different types of environments are more or less likely to have regular exposure to literacy activities, and early differences in experience are often linked to both brain development and educational outcomes (e.g., Burris et al., 2019; Noble et al., 2007; Romeo et al., 2022). Therefore, we also conducted secondary exploratory analyses examining associations between toddlers’ experience with reading activities and patterns of neural activity.

## Methods

### Participants

Forty participants were recruited from the community surrounding the campus of Tel Aviv University. This sample size was chosen to be larger than typical studies using fNIRS, accounting for a relatively high rate of attrition (Baek et al., 2022). The final sample included 31 monolingual, typically developing toddlers from the communities near the university campus (18 girls, 13 boys), ranging in age from 2.1-3.6 years (*M* = 2.8, *SD* = 0.44). Nine additional participants were tested but their data were excluded due to technical errors (3), providing too few channels of usable data after the preprocessing stage (5), or a reported diagnosis of autism (1). All children were tested using the text of their native language as reported by parents (Hebrew: *N* = 26, Russian: *N* = 3, English: *N* = 2), and none were enrolled in formal preschools or were taught reading. Parents provided written informed consent prior to participation, and families received a children’s book, a t-shirt and 40 NIS for their participation. Toddlers’ assent was assessed by monitoring their mood and verbality, and the experiment was terminated if the toddler showed or expressed discomfort. All procedures were approved by the Tel Aviv University IRB.

### Stimuli

Stimuli consisted of 50 written sentences of real text and 50 matched sequences of non-text symbols. In the Text condition, participants saw three-word child-directed sentences (e.g., “מהר רץ הכלב” [*The dog runs fast*], see Fig. 1A) presented in Courier New font. In the Non-text condition, participants saw 3-item sequences made up of nonsense characters in BACS-2 Sans font (Vidal et al., 2017). These characters are designed to share visual properties with text, but are unfamiliar and difficult to label (see Fig. 1B). Items in the Non-text condition were matched in length and number of unique characters to items in the Text condition. All sentences were presented one at a time in white text on black backgrounds. The text size corresponded to approximately 1.6° - 2.1° of visual angle vertically. The stimuli spanned between 15° −23° of visual angle horizontally. Each participant saw a randomly-sampled set of 20 sentences from each condition up to four times each.

Each trial included four sentences from a single condition (Text or Non-text). Sentences were matched across conditions for length and number of unique characters, and matched items were presented in a pseudo-random order within trials. No sentence was repeated within a block, and across participants, items were counterbalanced so that each stimulus appeared equally in the two conditions. Sentences were presented for 2 seconds. Each trial of 4 sentences was followed by an attention-getter (a colorful cartoon gif of a cat, dog, or sheep that played with fun instrumental music in the background). Attention-getters appeared in random order, each for a random duration of 8, 9, 10, 11 or 12 seconds to allow the hemodynamic response to return to baseline and to prevent the participant from being able to anticipate the onset of the next stimulus. Within a block, trials were randomized for each participant, with the constraint that each block contained an equal number of Text and Non-text trials, and so that no more than two consecutive trials were from the same condition.

### Home Literacy Questionnaire

Parents were asked to fill out the Home Literacy Environment Questionnaire (Van Steensel, 2006, translated to Hebrew, see OSF) prior to the experiment to provide information about their child’s exposure to activities related to reading (e.g., How often do you read bedtime stories to your child?; How often do you visit the library?). Questionnaire data was available for 29/31 participants. Parents’ responses on 13 questions were coded such that answers of “never” were scored as 0, “once or more a month” or “once or more a week” were scored as 1, and “every day” was scored as 2. We then calculated a sum (up to 26) for each participant (*M* =15.7/26, *SD* = 3.4) and divided participants into “High Literacy” (HLE > 15, M = 18.3, SD = 2.51, *N =* 15) and “Low Literacy” groups (HLE < 15, M = 13.1, SD = 2.05, *N* = 14), based on a median split. Dividing into groups enabled us to perform exploratory analyses testing whether toddlers with frequent exposure to literacy activities showed different patterns of neural activity than those with less frequent exposure. We chose this approach over testing correlations between individual scores and fNIRS parameters because individual GLM or time-window approaches assume a canonical hemodynamic response function, which is not widely accepted or available for this age group. In addition, our key manipulation involved subtle visual stimuli – relatively small black letters presented in silence – compared to the more engaging colorful attention getter paired with sound. Indeed, an exploratory GLM analysis yielded mostly negative beta values (see Table S1 in the Supplementary Materials).

### Procedure

Toddlers sat on their parents’ laps and watched stimuli on a 43” screen positioned 1.5 meters in front of them while wearing the fNIRS cap. Lights were turned off in the room to allow for better signal unless a participant expressed discomfort and the parent requested otherwise. Parents were asked not to direct their children’s attention to a specific type of sentences and not to read sentences aloud prior to or during the experiment and were not told the objective of the experiment until after they completed it.

Trials were presented in blocks of 10 trials. After each block, participants were allowed to take a break for as long as they wished. Overall, a participant that completed the entirety of the experiment saw 40 four-sentence trials, for a total of 80 Text sentences and 80 Non-text sentences. The experiment’s design is depicted in Fig. 1.

### Data Collection

Toddlers’ cortical responses were recorded with functional near-infrared spectroscopy (NIRSport2, NIRx). Toddlers wore a cap during the experiment to measure the changes of light properties and estimate the hemodynamic activity of their cortices. There were two cap sizes: 52 or 54 cm, measured and used depending on the toddlers’ age and head size. Each cap had 16 double tip sources (red dots in Fig. 2A), and 16 Silicon photodiode double detectors (blue dots in Fig. 2A). Overall, 48 channels were recorded, 24 in each hemisphere. The channels were divided such that 14 were in VLPFC, 12 in the STC, 10 in the pMD, and 12 in the Parietal cortex; divided equally between the Left and Right hemispheres in each region respectively. Additionally, we placed an accelerometer in the Pz location on the cap. Anatomical localization for channels is provided in a channel-wise mapping of fNIRS optodes to standardized MNI coordinates (Table 1). Each channel was assigned to cortical regions of interest (VLPFC, STC, pMD, and parietal cortex) based on its nearest correspondence to the MNI atlas used in Satori 2.2 (Brain Innovation, NIRx). The fNIRS cap was carefully centered on each toddler’s head and aligned so that the lower edge of the cap was positioned approximately 2 cm above the eyebrows, and cap placement was visually inspected and adjusted as needed to maintain symmetry and consistency across participants.

**Figure 2.**
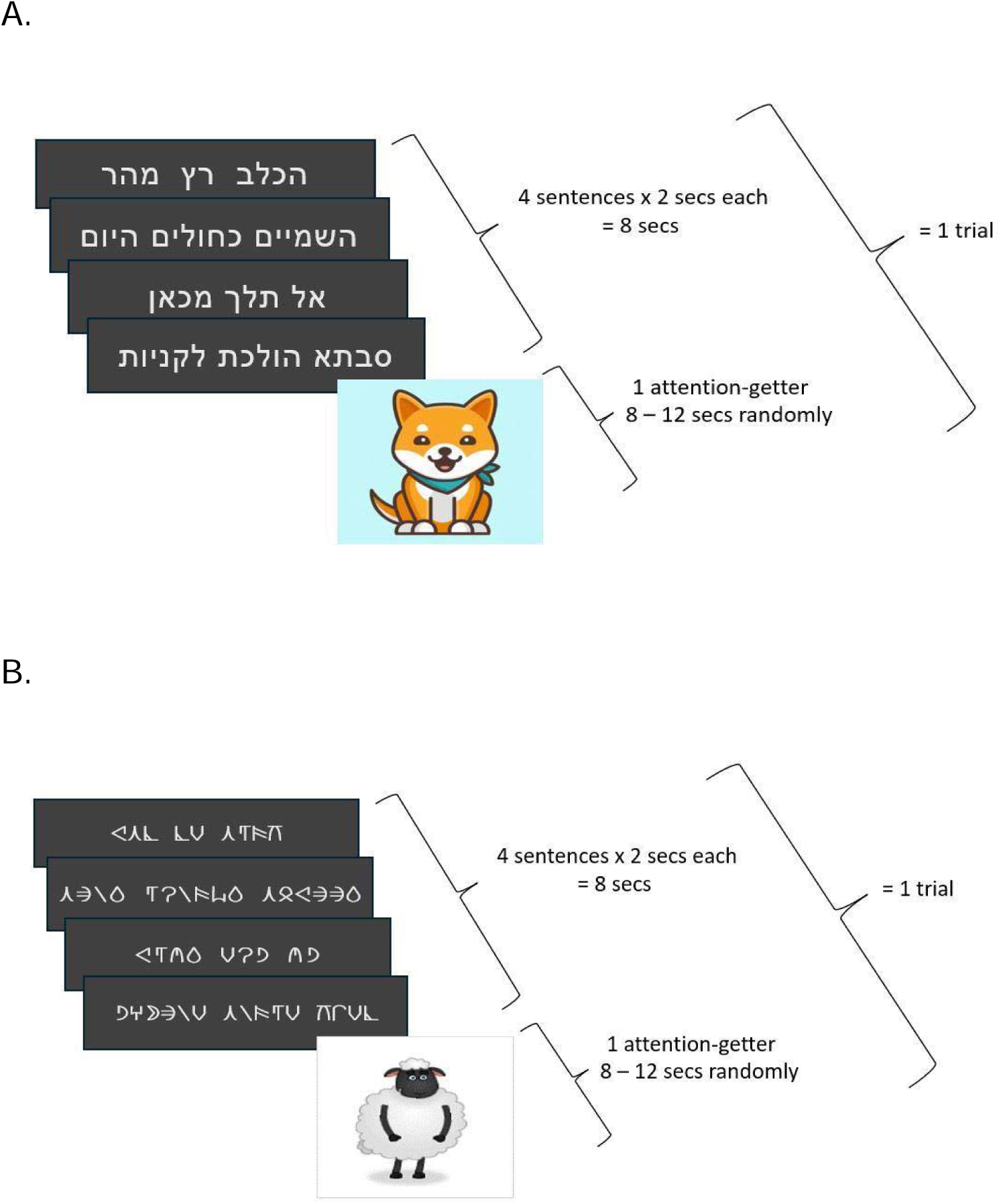
*Note*. Experimental design. A trial consisted of four consecutive sentences from the same condition each appearing for 2 seconds, followed by an animated attention-getter that appeared for a random duration of 8 - 12 seconds. There were up to 20 trials for each condition. **A**. Example of a trial in the Text condition (Hebrew). **B**. Example of a trial in the Non-text condition (BACS2 Sans font, adapted from Vidal et al., 2017).

**Figure 3.**
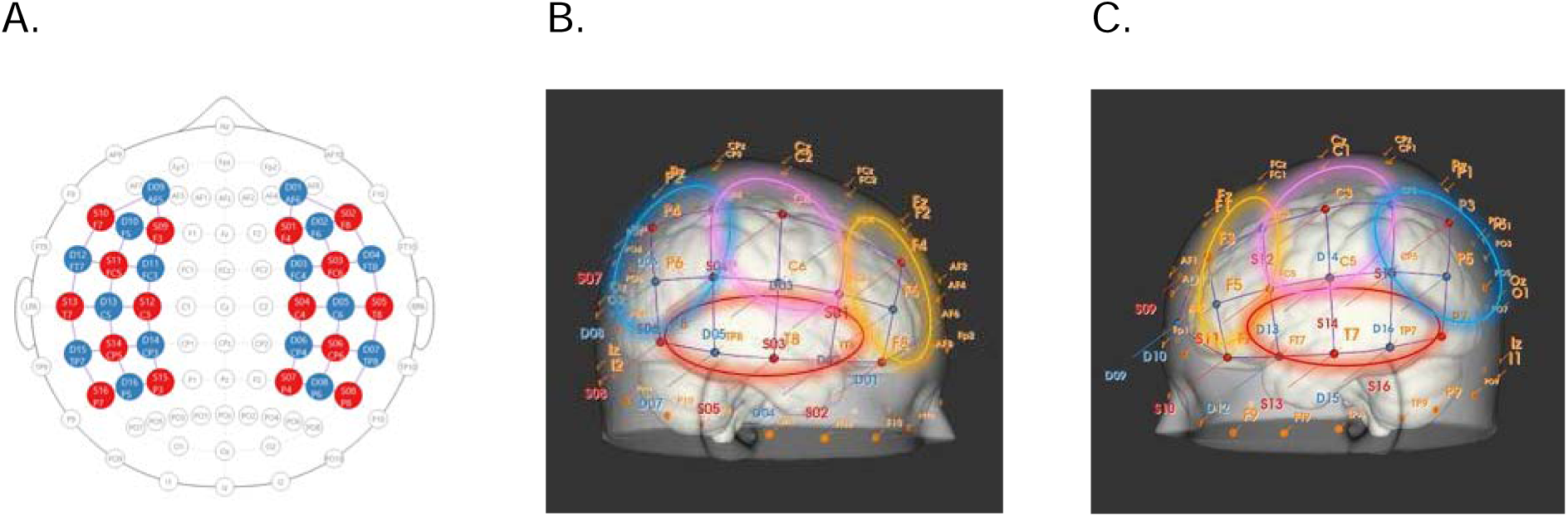
*Note*. fNIRS Cap Montage. **A**. 2D configuration: Red dots-sources. Blue dots-detectors. **B**. 3D Right view. **C**. 3D Left view. Channels in the ventral prefrontal cortex are highlighted in yellow. Channels in the superior temporal cortex are highlighted in red. Channels in the dorsal premotor cortex are highlighted in pink. Channels in the parietal cortex are highlighted in blue.

**Table 1.**

| ROI | Channel | MNI X | MNI Y | MNI Z |
| --- | --- | --- | --- | --- |
| Left VLPFC | S9-D9 | -39.5 | 52.2 | 15.9 |
|  | S9-D10 | -43.8 | 38.3 | 20.9 |
|  | S9-D11 | -44.8 | 25.2 | 38.4 |
|  | S10-D9 | -40.0 | 47.4 | -12.0 |
|  | S10-D10 | -53.8 | 35.4 | -5.3 |
|  | S10-D12 | -53.8 | 13.9 | -17.0 |
|  | S11-D10 | -57.9 | 22.9 | 14.6 |
| Right VLPFC | S1-D1 | 36.3 | 52.0 | 16.5 |
|  | S1-D2 | 42.8 | 44.1 | 18.1 |
|  | S1-D3 | 42.8 | 27.8 | 36.1 |
|  | S2-D1 | 45.9 | 45.0 | 11.4 |
|  | S2-D2 | 45.9 | 36.9 | -1.3 |
|  | S2-D4 | 54.8 | 14.9 | -16.9 |
|  | S3-D2 | 56.0 | 26.2 | 14.2 |
| Left STC | S11-D12 | -61.3 | -1.0 | 5.4 |
|  | S13-D12 | -63.4 | -18.2 | -11.4 |
|  | S13-D13 | -55.8 | -34.3 | 0.9 |
|  | S13-D15 | -68.6 | -29.4 | -8.2 |
|  | S14-D15 | -61.7 | -46.4 | 17.9 |
|  | S16-D15 | -60.1 | -55.8 | -6.0 |
| Right STC | S3-D4 | 55.6 | 9.7 | 0.5 |
|  | S5-D4 | 65.2 | -9.2 | -19.1 |
|  | S5-D5 | 66.5 | -15.5 | 6.1 |
|  | S5-D7 | 67.5 | -27.5 | -9.9 |
|  | S6-D7 | 66.1 | -34.0 | 7.7 |
|  | S8-D7 | 62.7 | -50.7 | -11.2 |
| Left PMd | S11-D11 | -58.0 | 15.0 | 28.3 |
|  | S11–D13 | -63.7 | -4.8 | 16.5 |
|  | S12–D11 | -46.7 | -4.0 | 54.7 |
|  | S12–D13 | -60.3 | -23.5 | 37.9 |
|  | S12–D14 | -52.3 | -33.1 | 53.6 |
| Right PMd | S3–D3 | 55.2 | 17.2 | 29.7 |
|  | S3–D5 | 59.8 | 16.9 | 5.2 |
|  | S4–D3 | 41.9 | 8.2 | 42.2 |
|  | S4–D5 | 60.3 | -5.4 | 29.7 |
|  | S4–D6 | 54.8 | -31.9 | 49.0 |
| Left Parietal | S14–D13 | -59.6 | -36.9 | 16.0 |
|  | S14–D14 | -49.3 | -53.9 | 39.5 |
|  | S14–D16 | -61.6 | -51.7 | 21.1 |
|  | S15–D14 | -34.2 | -53.5 | 57.8 |
|  | S15–D16 | -44.4 | -71.0 | 38.8 |
|  | S16–D16 | -54.9 | -68.6 | 3.4 |
| Right Parietal | S6–D5 | 46.9 | -22.8 | 20.7 |
|  | S6–D6 | 59.4 | -37.7 | 38.6 |
|  | S6–D8 | 57.9 | -56.1 | 19.1 |
|  | S7–D6 | 45.9 | -50.4 | 49.5 |
|  | S7–D8 | 50.0 | -67.5 | 26.9 |
|  | S8–D8 | 50.0 | -56.5 | 7.9 |

*Note.* MNI Coordinates represent the estimated spatial location of each fNIRS channel as provided in MNI atlas in Satori 2.2 (Brain Innovation, NIRx). Channel labels correspond to source–detector pairs.

### Data Preprocessing

Participants’ recordings were included in the analysis only if they successfully completed at least four trials in each condition (almost one full block). The data was preprocessed with Satori software (Brain Innovation, NIRx). Raw voltages were used for channel rejection with coefficient variation (CV) threshold set at 45%. Voltages were transformed into optical density (OD). Spikes were removed from OD with the following set parameters: iterations at 10, lags at 5 seconds, threshold at 3.5, influence at 0.5, and interpolation set to monotonic. Interpolated OD was bandpass filtered using a Butterworth filter between 0.01 and 0.5 Hz. Filtered OD was transformed into chromophore concentrations (CC). Finally, CC was normalized via z-transformation.

### Data Analysis

Event-related averages (ERAs) of oxyhemoglobin (HbO_2_) changes were extracted between −5 and 25 seconds from stimulus onsets and averaged for each participant per region of interest (ROI). The 48 channels were divided into four ROIs: ventral prefrontal cortex, superior temporal cortex, dorsal premotor cortex, and parietal cortex. The time window between −5 to 0 secs was used for baseline correction. An ROI was calculated if there were at least two channels left after preprocessing. On average, after preprocessing, there were 6.4 usable channels in the Left VLPFC (8.6% rejected), 6.7 channels in the Right VLPFC (4.3% rejected), 5.4 channels in the Left STC (10% rejected), 5.6 channels in the Right STC (6.7% rejected), 4.6 channels in the Left pMD (8% rejected), 4.6 channels in the Right pMD (8% rejected), 5.3 channels in the Left Parietal (11.7% rejected), and 5.1 channels in the Right Parietal (15% rejected). On average, toddlers completed 25.5 trials with 12.9 trials for the text condition, and 12.6 trials for the non-text condition. Participants with fewer than four usable trials per condition were excluded from analyses. A two-tailed paired t-test was used to test if the difference between the Text and Non-text conditions was significant across participants for each timepoint, with a cluster-size correction method (*p* < 0.05; Maris & Oostenveld, 2007). Since different channels were rejected for each toddler, we focused our main analyses on ERA by ROIs, which allowed us to keep the same degrees of freedom in all the statistical tests with greater reliability within each toddler, while allowing for flexibility regarding the dynamics of the expected response.

## Results

Data was analyzed using Python. All data, materials, and processing scripts are available through OSF [https://osf.io/zswgx/overview?view_only=0f6911de4e7d4a588790e05733a6ea95].

### Comparing cortical activity for Text vs. Non-text

The oxygenated Hemoglobin levels differed between the two conditions in toddlers’ cortices. The response in the left VLPFC was stronger for the Text condition than for the Non-text condition; a significant difference was found at 6.5 - 12 seconds post stimulus onset (*t*(30) = 2.36, *p* = .02, with a paired-samples Hedges’ *g* = .41, 95% CI [.08, .89]) (Fig. 4A). On the other hand, no difference was observed in the right VLPFC (*t*(30) = - .55, *p* = .58, with paired-samples Hedges’ *g* = -.09, 95% CI [-.49, .24]) (Fig. 4B). Furthermore, the response in the STC was stronger for the Non-text condition than for the Text condition; a significant difference in the left STC was found 3.4 - 20.4 seconds post stimulus onset (*t*(30) = −3.29, *p* = .002, with a paired-samples Hedges’ *g* = -.57, 95% CI [-.98, -.27]) (Fig. 4C), and a trend in the right STC 9.5 - 12.5 seconds post stimulus onset (*t*(30) = −1.79, *p* = .08, with a paired-samples Hedges’ *g* = -.31, 95% CI [- .78, .03]) (Fig. 4D). Meanwhile, no difference was observed in the left pMD (*t*(30) = -.59, *p* = .55, with a paired-samples Hedges’ *g* = -.1, 95% CI [-.45, .26]), or the right pMD (*t*(30) = −1.18, *p* = .24, with a paired-samples Hedges’ *g* = -.2, 95% CI [-.58, .13]) (Fig. S1A, S1B). Lastly, no difference was observed in the left parietal cortex (*t*(30) = 1.08, *p* = .28, with a paired-samples Hedges’ *g* = .19, 95% CI [-.15, .52]), or the right parietal cortex (*t*(30) = -.04, *p* = .96, with a paired-samples Hedges’ *g* = -.008, 95% CI [-.48, .28]) (Fig. S1C, S1D). To confirm that effects were consistent across languages, an analysis including only Hebrew-speaking participants (*N* = 26) revealed similar patterns (see Fig. S2 on OSF). Deoxygenated Hemoglobin (HbR) levels between the two conditions significantly differed in the pMD. In the left pMD, HbR levels were stronger for the non-text condition than the text condition at 15.6 - 25 sec post stimulus onset. Meanwhile, in the right pMD, HbR levels were stronger for the non-text condition than the text condition at 13 – 25 sec post stimulus onsets. Aside from that, HbR levels did not differ between the two conditions in any of the other ROIs. ERAs analysis for deoxygenated Hemoglobin levels (HbR) can be found on OSF (Fig. S3).

**Figure 4.**
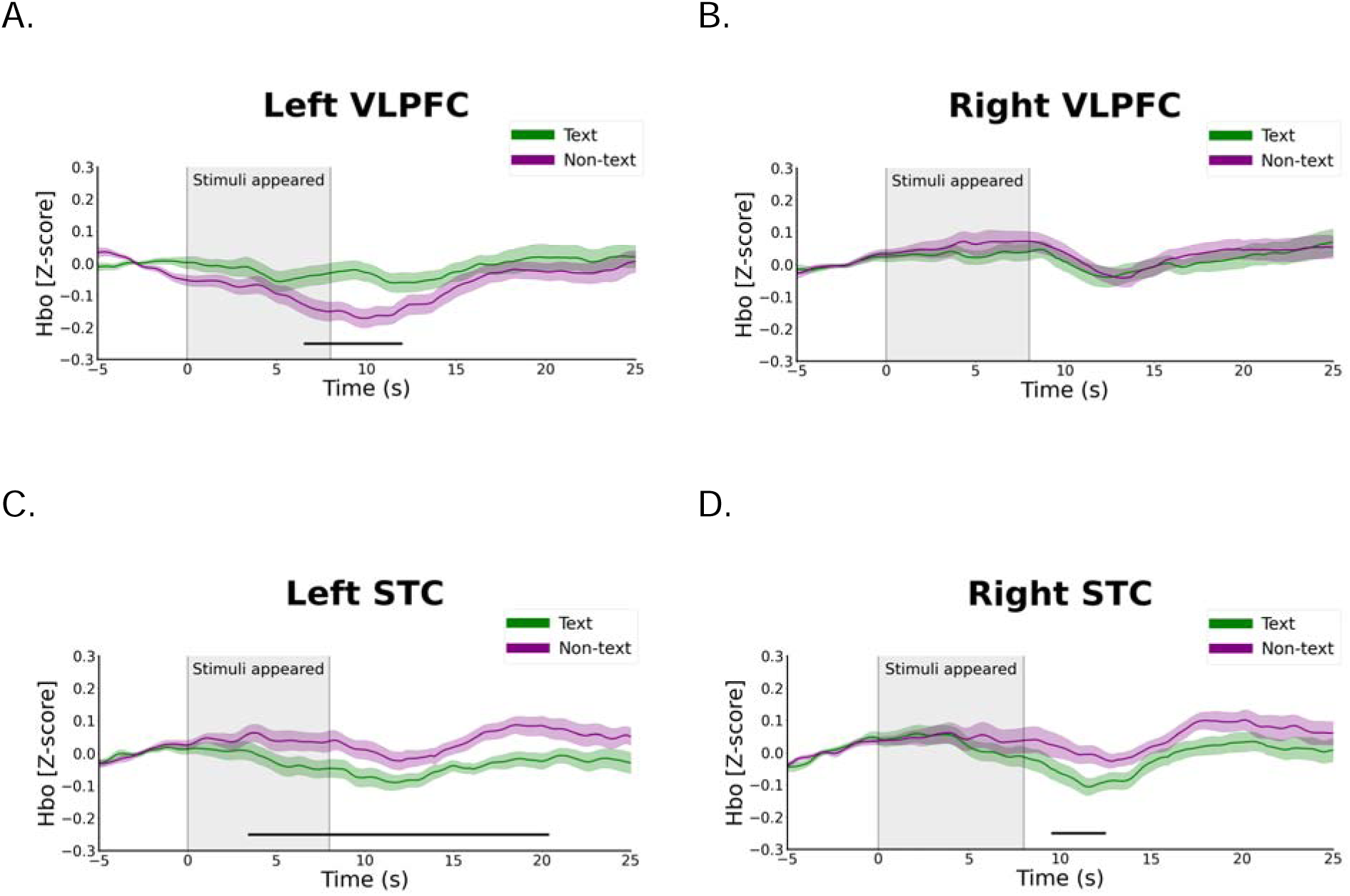
*Note.* Event-related averages of HBO_2_ by cortical ROI. Green–Text condition. Purple–Non-text condition. Shaded patches represent Standard Errors. Black bars under each graph denote the time windows where the difference between the conditions was significant (*p* < .05). **A**. Left ventrolateral prefrontal cortex. Text > Non-text at 6.5 - 12 sec post stimulus onset. **B**. Right ventrolateral prefrontal cortex. **C**. Left superior temporal cortex. Non-text > Text at 3.4 - 20.4 sec post stimulus onset. **D**. Right superior temporal cortex. Non-text > Text at 9.5 – 12.5 sec post stimulus onset.

### Lateralization of response to text

The condition difference in oxygenated Hemoglobin levels differed between toddlers’ left and right cortices, suggesting signs of lateralization. The cortical activity was stronger for the right VLPFC than for the left VLPFC in the Text condition at 3.5 - 9.2 sec post stimulus onset (t(30) = −1.69, p = .05, with a paired-samples Hedges’ g = -.29, 95% CI [- .67, .04]), as well as in the Non-text condition at 0 - 25 sec post stimulus onset (t(30) = - 5.28,p < .001, with a paired-samples Hedges’ g = -.92, 95% CI [-1.54, -.55]). Furthermore, the cortical activity was stronger for the right STC than for the left STC in the Text condition at 0 - 7 sec post stimulus onset (t(30) = −1.97, p = .03, with a paired-samples Hedges’ g = -.34, 95% CI[-.69, -.02]), but not in the Non-text condition (p > .05 across all time points; Fig. S4D). No differences emerged between the left- and right-pMD or Parietal cortex in either condition (p > .05 across all time points; Fig. S4E-H). While results suggest that the cortical activity differs in response to both Text and Non-text conditions, in the VLPFC and the STC, differences were more prominent in the left hemisphere. Overall, the responses in the right cortices were stronger than in the left cortices, but the responses in left cortices were more sensitive to the nature of the presented stimuli. Figures depicting detailed comparisons between left and right ROIs can be found on OSF (Fig. S4).

### Exploratory associations between text exposure and cortical activity

The oxygenated Hemoglobin levels differed in the High Literacy group and Low Literacy group in both right and left PMd for both Text and Non-text conditions, and in the right VLPFC and left Parietal ROIs in the Non-text condition (see Figure S8 in the supplementary for more detailed description of the HLE effects. Additionally, cortical responses differed between the Text vs. Non-text conditions, across the High Literacy group (High HLE) and the Low Literacy group (Low HLE) in multiple ROIs:

#### High Literacy Group

The cortical response was stronger for the Text condition than for the Non-text condition in the left VLPFC; a marginally significant difference was found at 8.3 - 11.8 sec post stimulus onset (*t*(14) = 1.77, *p* = .09, with a paired-samples Hedges’ *g* = .43, 95% CI [-.03, 1.38]), while no difference was found in the right VLPFC (*t*(14) = .06, *p* = .55, with a paired-samples Hedges’ *g* = .14, 95% CI [-.37, .7]). Moreover, no difference was found in the left STC (*t*(14) = -.92, *p* = .36, with a paired-samples Hedges’ *g* = -.22, 95% CI [-.74, .3]), while the cortical response was stronger for the Non-text condition than for the Text condition in the right STC at 11.3 - 12 sec post stimulus onset (*t*(14) = -.7, *p* = .49, with a paired-samples Hedges’ *g* = -.17, 95% CI [- .91, -.31]). The cortical response was stronger for the Non-text condition than for the Text condition in the left pMD (*t*(14) = 1.09, *p* = .29, with a paired-samples Hedges’ *g* = .26, 95% CI [-.22, 1.02]), while no difference was found in the right pMD (*t*(14) = .91, *p* = .37, with a paired-samples Hedges’ *g* = .22, 95% CI [-.32 .68]). Lastly, no differences were found between the conditions in the parietal cortex (all p > .1, all 95% CI include 0).

#### Low Literacy Group

No difference was found in either the left VLPFC (*t*(13) = .13, *p* = .89, with a paired-samples Hedges’ *g* = .03, 95% CI [-.62, .55]), or the right VLPFC (*t*(13) = −1.1, *p* = .29, with a paired-samples Hedges’ *g* = -.27, 95% CI [-.92, .22]). The cortical response was stronger for the Non-text condition than for the Text condition in the left STC; a significant difference was found at 8.1 - 9.7 and 13.1 - 20.8 sec post stimulus onset (*t*(13) = −3.02, *p* = .009, with a paired-samples Hedges’ *g* = -.76, 95% CI [-1.44, - .36]), and in the right STC at 17.3 - 18.2 sec post stimulus onset (*t*(13) = −1.37, *p* = .19, with a paired-samples Hedges’ *g* = -.34, 95% CI [-1.06, .12]). Moreover, the cortical response was stronger for the Non-text condition than for the Text condition in the left pMD at 11.3 - 22.2 sec post stimulus onset (*t*(13) = −2.22, *p* = .04, with a paired-samples Hedges’ *g* = -.56, 95% CI [-1.13, -.14]), and in the right pMD at 13.5 - 14.1 sec post stimulus onset (*t*(13) = −2.41, *p* = .03, with a paired-samples Hedges’ *g* = -.61, 95% CI [-1.22, -.28]). Lastly, no differences were found between the conditions in the Parietal cortex (all p > .1, all 95% CI include 0). ERAs analysis for oxygenated Hemoglobin levels for high HLE and low HLE can be found on OSF (Fig. S5).

### Exploratory associations between age and cortical activity

Participants were divided into two groups based on the median split of age (*Med* = 2.7 years; Younger toddlers: *N* = 15, *M* = 2.4 years, *SD* = 0.15; Older toddlers: *N* = 16, *M* = 3.1 years, *SD* = 0.28). The two age groups exhibited similar responses to both conditions in all ROIs and similar sensitivity to the Text condition (vs. Non-text condition) in the left VLPFC, but showed different pattern of responses in STC, where the older group showed a novelty response to Non-text (vs. Text condition).

#### Younger toddlers

On average, younger toddlers completed 25.7 trials, with 12.5 trials included for the Text condition, and 13.2 were included for the Non-text condition. On average, after preprocessing, there were 6.7 usable channels in the Left VLPFC, 6.7 channels in the Right VLPFC, 5.4 channels in the Left STC, 5.5 channels in the Right STC, 4.7 channels in the Left pMD, 4.5 channels in the Right pMD, 5.3 channels in the Left Parietal, and 4.7 channels in the Right Parietal.

The cortical response was stronger for the Text condition than for the Non-text condition in the left VLPFC; a significant difference was found at 7.5 - 10 seconds post stimuli onset (*t*(14) = -.79, *p* = .09, with a paired-samples Hedges’ *g* = -.4, 95% CI [-1.03, .01]), whereas no difference was found in the right VLPFC (*t*(14) = .03, *p* = .97, with a paired-samples Hedges’ *g* = -.16, 95% CI [-0.88, .3]). No difference was found in either the left STC (*t*(14) = −1.79, *p* = .09, with a paired-samples Hedges’ *g* = -.43, 95% CI [-1.03, .01]), or in the right STC (*t*(14) = .07, *p* = .93, with a paired-samples Hedges’ *g* = .01, 95% CI [-.05, .57]). Moreover, no differences were found between the conditions in the pMD or the Parietal cortex (all p > .1, all 95% CI include 0).

#### Older toddlers

On average, older toddlers completed 25.3 trials, which included 13.2 tials for the text condition, and 12.1 trials for the non-text condition. On average, after preprocessing, there were 6.6 usable channels in the Left VLPFC, 6.6 channels in the right VLPFC, 5.3 channels in the Left STC, 5.6 channels in the right STC, 4.7 channels in the left pMD, 4.6 channels in the right pMD, 5.5 channels in the left Parietal, and 5.5 channels in the right Parietal.

The cortical response was stronger for the Text condition than for the Non-text condition in the left VLPFC; a significant difference was found at 8.9 - 11.3 seconds post stimuli onset (*t*(15) = 1.63, *p* = .12, with a paired-samples Hedges’ *g* = .38, 95% CI [-.08, 1.2]), whereas no difference was found in the right VLPFC (*t*(15) = -.68, *p* = .5, with a paired-samples Hedges’ *g* = .38, 95% CI [-.08, 1.2]). Moreover, the cortical response was stronger for the Non-text than for the Text condition in the STC; a significant difference was found at 1 - 7.5 seconds post stimuli onset in the left STC (*t*(15) = −2.85, *p* = .01, with a paired-samples Hedges’ *g* = -.67, 95% CI [-1.36, -.28]), and at 9.8 - 25 seconds post stimuli onset, and in the right STC (*t*(15) = −2.73, *p* = .01, with a paired-samples Hedges’ *g* = -.65, 95% CI [-1.55, -.16]). Furthermore, no differences were found between the conditions in the pMD or the Parietal cortex (all p > .1, all 95% CI include 0). ERAs analysis for oxygenated Hemoglobin levels for younger toddlers and older toddlers can be found on OSF (Fig. S6).

## Discussion

This study provides the first evidence that neural regions implicated in reading show responsiveness to text well before children can read. Previous research emphasizes the slow development of the reading network, suggesting cortical specialization emerges only after children have extensive practice and achieve relatively high skill (Cantlon et al., 2017; Liu et al., 2018; Schlaggar & McCandliss, 2007; Smith et al., 2018). In contrast, we found that toddlers with no formal schooling showed different patterns of brain activity in response to text than to equally complex non-text symbols, suggesting text was treated as a familiar stimulus. Strikingly, we found a left-lateralized response to text in the ventrolateral prefrontal cortex, which could reflect that toddlers have begun to detect associations between text and language. Based on these results, we suggest infants’ implicit attention to patterns in their environments (Saffran & Kirkham, 2018) could form the foundation for their later ability to build connections between representations of a word’s meaning, sound, and visual form that enable them to become fluent readers (see Seidenberg, 2017).

The finding that text evoked different neural activity than non-text symbols across multiple cortical regions supports our proposal that pre-reading toddlers have identified text as a meaningful stimulus. Notably, toddlers showed heightened activity in response to text in the left VLPFC, but not the right VLPFC. The left VLPFC has long been implicated in processing language and in selecting and retrieving information from semantic memory (Badre & Wagner, 2007). Given that toddlers viewed text in silence, this selectivity suggests they may have begun to connect text with language before they can decode the visual signal. Studies with five-year-old pre-reading children suggest they show a left-lateralized response to text in visual regions that is correlated with behavioral measures of pre-reading (Lochy et al., 2016), providing additional evidence that lateralization may precede skilled reading. Here, we found evidence of this lateralization in children several years younger than in prior studies and in language rather than visual regions, suggesting toddlers could be learning about the function as well as the form of text.

While our results could reflect that children may be beginning to form associations between text and language, it is important to acknowledge that left VLPFC activity could also reflect non-linguistic processes, such as working memory or attentional control (Badre & Wagner, 2007; Baüml et al., 2010; Weintraub-Brevda et al., 2019). We did find that children who participated more in literacy activities showed more reliable differences in responses to text vs. non-text in the left VLPFC, consistent with the idea that children build these associations through experience. However, it is possible that children who have more experience with reading activities are building other domain-general skills, and these effects are not directly attributable to their familiarity with the linguistic function of text. For example, it could be that children who engage in shared reading more often have different attentional strategies, and those attentional differences lead them to respond more strongly to a familiar stimulus. Future research will be needed to better understand why text elicited a familiarity response in left VLPFC, but the current results do provide evidence that text evokes a different response in this region than in other regions that we measured. Interestingly, this effect was already apparent in younger toddlers in our sample, suggesting this could be one of the earliest emerging regions in the reading network

Activity in other regions also supports the view that toddlers recognize text as familiar. In both left and right STC, toddlers’ brains showed more activity in response to unfamiliar symbols. While this increased novelty response did not align with our predictions, research using other neuroimaging methods has found that novel stimuli elicit activity in the superior temporal gyrus (Optiz et al., 1993). Thus, toddlers appear to have interpreted non-text symbols as novel, which suggests they treated text as familiar. This early differentiation contrasts with results showing that different types of symbols (letters, digits) yield activity in similar regions for children not yet reading or just beginning to learn (Cantlon et al., 2011; Crollen et al., 2025). Our use of symbols that toddlers had never encountered may have made it easier to detect toddlers’ growing familiarity with text, perhaps before they have well-established representations of letters. On the other hand, we did not observe differences across conditions in the parietal cortex. Parietal activity has been linked to letter-by-letter reading in early stages of literacy, when children may be working to form connections between visual forms and sounds (Dehaene et al., 2010; Moulton et al., 2019), and pre-reading toddlers may not yet separate individual letters and are unlikely to have learned letter-sound pairings. Deoxygenated Hemoglobin (HbR) recordings in the dorsal premotor region (left and right PMd) also show heightened responses to visually presented text over non-text, suggesting involvement of a possible articulation component in distinguishing between these two conditions (Dehaene, 2009) and providing another illustration that these two types of stimuli were processed differently. It should also be acknowledged that using fNIRS, we could not measure activity in occipital areas including the VWFA, limiting our ability to draw conclusions about the role of visual regions. Future research will be needed to map specific knowledge and behaviors associated with reading onto relevant neural correlates, but our study offers new insight into the early stages when parts of what will become the reading network start to develop preferential responses to text.

Improvements in reading are associated with specialized neural processing, but it is not yet known whether early specialization would predict later skill for individual children. Children with reading difficulties show different patterns of neural activity and lateralization than peers with stronger reading skills (Banfi et al., 2019; Hoeft et al., 2007; Norton et al., 2015). We do not have behavioral measures to connect children’s brain activity and pre-reading skills either concurrently or over time, but future research could test associations between experience with reading, visual and oral pre-reading skills, and reading outcomes. It would also be informative to test whether these regions develop similarly for children learning to read in different languages. While we found no evidence that including participants who were not learning Hebrew affected our results, future research could more systematically test whether there are differences for children exposed to writing systems with shallower or deeper orthography, for example, or those learning character-based scripts vs. alphabetic systems (see Zhang et al., 2024). Understanding how neural organization co-develops with behavioral changes will allow us to better explain the emergence of expertise and could offer tools for identifying and supporting children at risk.

Engaging in literacy activities supports children’s reading skills, and explanations tend to emphasize how oral language and reading skills are mutually reinforcing (Farrant & Zubrick, 2012; Mol & Neuman, 2014). While reading experiences undoubtedly offer valuable opportunities for learning language, children may also gain familiarity with the visual form of language and begin establishing the neural circuitry that will allow them to read efficiently. Our results show that years before children reliably know the letters of the alphabet, let alone read fluently (Share & Blum, 2005), they have begun to process text differently than other stimuli. These patterns of activity illustrate a potential pathway through which the reading network may develop. Infants’ well-established statistical learning abilities could allow them to detect the frequent presence of text in their environments and its co-occurrence with rich language input and in turn, lead them to begin to build neural connections between language and reading-related regions. Ultimately, we suggest that changes in the brain associated with reading appear in very early phases of the learning process, before the appearance of obvious behavioral signals (fluent reading or experience with formal instruction), and not as a subsequent response to the acquisition of a new skill.

## Acknowledgements

We would like to thank the research team at the TAU Center for Cognitive Development, especially Helen Antipko, Salma Kabaha, Liliya Kristalev and Yarin Zarfati.

